# SoftAlign: End-to-end protein structures alignment

**DOI:** 10.1101/2025.05.09.653096

**Authors:** Jeanne Trinquier, Samantha Petti, Sukhwan Park, Kithmini Herath, Michel van Kempen, Shihao Feng, Johannes Söding, Martin Steinegger, Sergey Ovchinnikov

## Abstract

With the recent breakthrough of highly accurate structure prediction methods, there has been a rapid growth of available protein structures. Efficient methods are needed to infer structural similarity within these datasets. We present an end-to-end alignment method, called SoftAlign, which takes as input the 3D coordinates of a protein pair and outputs a structural alignment. In addition to the traditional Smith-Waterman alignment method, we introduce a modified softmax alignment that shows very promising results for structure similarity detection. We demonstrate that the SoftAlign model is able to recapitulate TM-align alignments while running faster, and it is more accurate than Foldseek on alignment and classification tasks. Although SoftAlign is not the fastest method available, it is highly precise and can be used effectively with other prefilters. In addition to developing an end-to-end structural aligner, our main contribution is the introduction and analysis of a pseudo-alignment method based on softmax, which can be used with other architectures, even those not based on structural information. The code for SoftAlign is available at https://github.com/jtrinquier/SoftAlign.

---

With the recent success of methods such as AlphaFold2 [1] and RoseTTAFold [2], hundreds of millions of protein structures with near-experimental quality are available. This offers a new horizon for understanding how to relate sequence, structure, and function. One application of structure comparison is the improvement of homology search, the task of identifying similar regions in protein sequences likely due to shared ancestry. Biologists use homology search to build databases of annotated proteins [3] and formulate hypothesis by comparison. Given a query protein with an unknown biological function, a common strategy is to use sequence similarity search in order to find homologous sequences with a known function [4, 5]. Homology search methods based on sequence similiarity have been very powerful and reliable when the query and the homologs exhibit a high sequence similarity. However, it is still a challenge to annotate proteins using just the sequence information when there is a low sequence similarity [6]. Incorporating structure comparison into homology search offers the potential to improve the detection of remote homologs when structure has been conserved. Indeed, alignment methods using structural information have been shown to offer a higher sensitivity at longer evolutionary distance [7]. It is important to note, however, that structural similarity does not necessarily imply homology—similar structures may arise independently through convergent evolution. Nevertheless, structural similarity can still provide valuable insights, such as identifying shared functional mechanisms, suggesting similar biochemical roles, or classifying proteins into common folds or architectures, even in the absence of sequence homology. Standard state-of-the art structural alignment tools such as TM-align [8] and Dali [9] are too computationally expensive to apply to a large scale comparison of structures in these huge new databases of protein structures. The first reason is that sequence based search tools use fast pre-filter algorithms that speed up the search by orders of magnitude. Second, the structural similarity scores computed by these methods are non-local, meaning that efficient dynamic programming algorithms such as Smith-Waterman cannot be used.

Our model, SoftAlign, takes as inputs the 3D coordinates of each protein structure, then transforms the coordinates of each position into a vector of features using a retrained encoder of the ProteinMPNN neural network [10]. A similarity matrix is obtained by taking the the scalar products of the vectors at all pairs of positions between the two proteins. Under the assumption that the feature vectors encode relevant information about the structure, we expect pairs of positions that should be aligned to have similar feature vectors, and thus to have a high scalar product.

To create an alignment from this similarity matrix, we tested two strategies. The first one, that we already partially developed in [11], is to apply a differentiable and smooth version of Smith-Waterman to [12] this similarity matrix to obtain an alignment. Using a differentiable version of the Smith-Waterman algorithm allows us to include the alignment step when training the neural network, making our approach end-to-end. The second approach is to apply a modified softmax function directly to the similarity matrix. This approach, while creating a pseudo alignment, demonstrates superior sensitivity for structure search compared to traditional methods. Notably, it achieves state-of-the-art results on a benchmark task with structures from SCOPe40 dataset [13].

We also present a categorical version of the network, where the structure is embedded into categorical variables. This approach is inspired by Foldseek [14], a structure comparison method that discretizes both query and target protein structures into sequences over an alphabet of 20 letters, known as 3Di states. While this approach shows similar results as Foldseek that is using a similar categorical strategy, continuous embedding methods outperforms this categorical strategy by a huge margin. As noted in the Discussion Section, the limitation primarily comes from the inability of the softmax method to effectively function with the categorical version. Despite this drawback, we continue to present the categorical version because constructing an alphabet that incorporates structural information is crucial for advancing protein language development [15]. Nevertheless, our results indicate that, at present, there is limited utility in using our alphabet for pre-filtering compared to Foldseek. This aspect will be left for future exploration and refinement.

## Related Work

### Standard structural aligners

Structural alignment tools such as TM-align [8] or DALI [9] can align protein structures using superposition, even when sequence based methods fail because of low-sequence similarity. However, these algorithms do not use dynamic programming to determine superposition and are very computationally expensive. Therefore, they are not well adapted for databases with hundred of millions of structures.

### Foldseek

Our work builds on the Foldseek structure comparison method, which discretizes the query and target structures into sequences over an alphabet of 20 letters learned using a vector-quantized variational autoencoder [14]. The 20 states, called 3Di states describe for each residue the geometrical conformation with its closest neighbors. This transformation from the structure to a sequence enables use of existing sequence search tools [16] with highly optimized implementations, speeding up by orders of magnitude the search while giving similar results to structural alignment methods such as TM-align [8]. Unlike our model, the Foldseek network is not trained end-to-end; the 3Di states are learned by training an autoencoder, and then a BLOSUM-like substitution matrix is computed from the empirical substitution frequencies of the 3Di states.

### Protein language based methods

In recent years, there has been a proliferation of large protein language models (PLLMs) trained on extensive datasets [17]. These models generate embeddings that contain rich information about protein sequences. As a result, these embeddings can be used for various bioinformatics tasks. New methods use the similarity between embeddings to detect distant homologous relationships between proteins [18, 19, 20]. Moreover, these embeddings can be used to create similarity matrices through simple operations like scalar products. Then, alignment algorithms such as Smith-Waterman can be applied to create an alignment. It is interesting to note that our softmax pseudo-alignment method could be integrated into the workflow of these methods.

## Results

We present SoftAlign, an end-to-end network that generates soft alignments from a similarity matrix derived from 3D protein coordinates. This workflow is summarized in Figure 1. We compare two differentiable alignment strategies: the Smooth Smith-Waterman algorithm and the softmax-based alignment. The Smooth Smith-Waterman method, adapted from [12], replaces the max operation with logsumexp, producing a probability distribution over alignments and enabling gradient-based optimization. The softmax-based approach, on the other hand, applies row- and column-wise softmax transformations to the similarity matrix (see Equation 1 in the Methods section), resulting in a pseudo-alignment that does not enforce sequential consistency. While Smooth Smith-Waterman maintains structural constraints by preserving order, the softmax approach provides a more flexible but potentially less structured alignment. Figure 2 illustrates the differences between these two methods.

**Figure 1.**
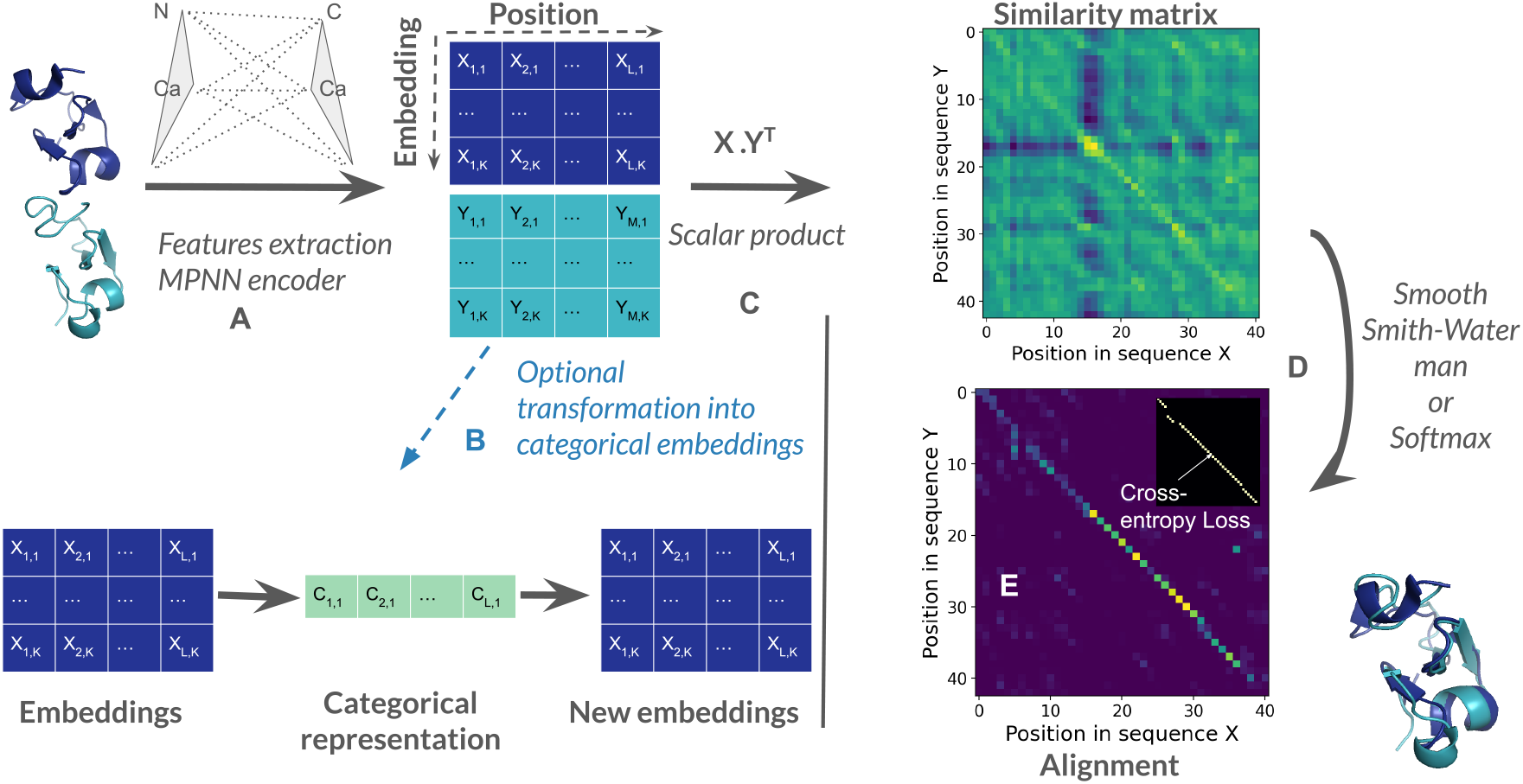
Network architecture overview. Starting from the 3D coordinates of a protein pair, the encoder extracts key structural features: interatomic distances, and backbone dihedral angles. These features are embedded using a retrained MPNN encoder [10], yielding an embedding matrix for each structure of size *L* × embedding dim (a). Optionally, embeddings can be converted into a categorical representation (b), as detailed in the Methods Section. A similarity matrix is derived by multiplying the embedding matrices (c), with each element (*i, j*) representing the alignment score between position *i* of the first protein and position *j* of the second. Soft alignment is achieved by applying either Smooth-Smith-Waterman or a softmax function to the similarity matrix (d). Training is guided by a cross-entropy loss, comparing the predicted alignment to a reference alignment (e).

**Figure 2.**
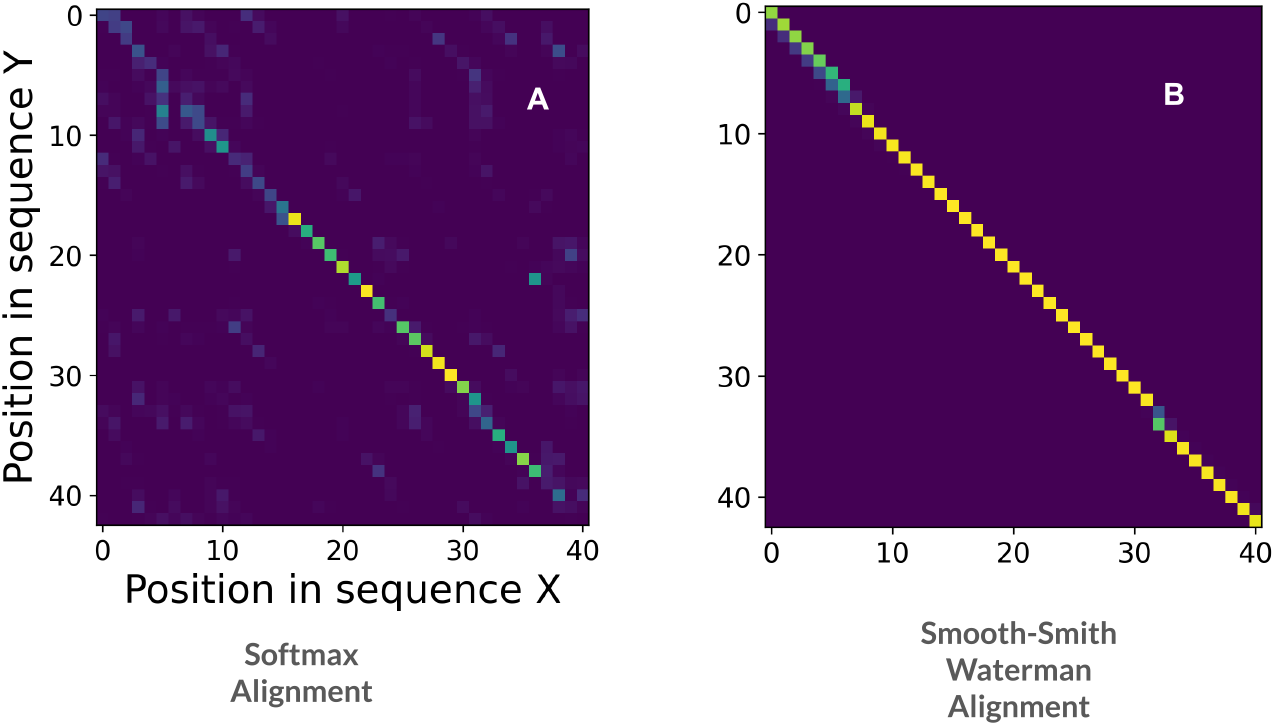
Comparison of alignment methods. The image in (a) represents the softmax alignment, while (b) illustrates the Smooth Smith-Waterman (SSW) alignment. The intensity reflects the score of aligning position i with j. Unlike SSW, the softmax alignment does not enforce a strict sequential structure—alignments can occur in arbitrary matrix positions. In contrast, SSW enforces sequential consistency, meaning alignment paths can only proceed down or to the right in the matrix, preserving the order of residues in both sequences.

We evaluated our method on two different tasks: pairwise alignment and structure classification based on structure similarity.

## Alignment quality

We evaluated the performance of our models on a test set of non-seen folds comprising 1000 pairs from SCOPe40 [13], using LDDT and TM-scores as metrics for alignment quality. These pairs were specifically selected to have a TM-score *>* 0.6, ensuring they represent biologically meaningful alignments, as TM-scores above this threshold generally indicate significant structural similarity. TM-align served as our reference point due to its highest level of alignment accuracy, thus setting a benchmark for comparison. Table 1 presents the alignment quality comparison. It is important to clarify the differences in training and inference configurations for the SoftAlign (Sfa) variants. The Sfa SW model is trained using the Smith-Waterman (SW) algorithm and also applies SW during inference. The Sfa Softmax variant, on the other hand, is trained and evaluated using a softmax-based pseudo-alignment approach, which does not enforce strict alignment paths. To address the limitations of this approximation, the Sfa Softmax rescued variant is trained with softmax but uses the Smith-Waterman algorithm during inference to refine the alignments, resulting in improved performance. The Sfa SW LDDT Loss model is trained with SW-based alignment, but instead of using standard cross-entropy loss, it employs a smoothed LDDT loss (see Equation 7 in the Methods Section). Additionally, the Categorical Sfa model, like the Sfa SW model, is trained with Smith-Waterman alignment and applies the same SW method during inference.

**Table 1:**
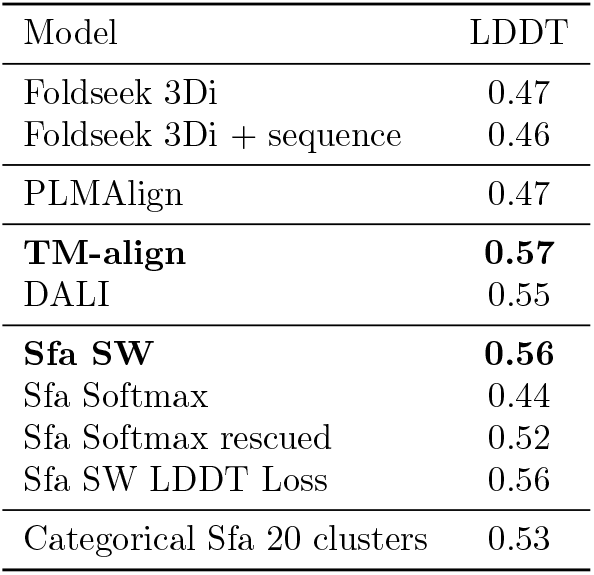
Comparison of mean LDDT scores across different alignment methods. TM-align and DALI serve as traditional structural alignment baselines, while Foldseek and PLMAlign represent alternative sequence- and structure-based approaches. Our SoftAlign (Sfa) models achieve competitive performance, with the Smith-Waterman SoftAlign (Sfa SW) reaching an LDDT score of 0.56, close to TM-align. The “rescued” variant of Sfa Softmax refers to applying the Smith-Waterman algorithm at inference time, improving its alignment quality from 0.44 to 0.52.

The Smith-Waterman SoftAlign (Sfa SW) model emerges as the second-best aligner for mean LDDT scores. In particular, its performance is comparable to that of standard aligners like TM-align and DALI and surpasses all other aligners by a big margin. Specifically, TM-align achieves a mean LDDT score of 0.57, while our SoftAlign model achieves a competitive score of 0.56 for LDDT, outperforming Foldseek and other baselines.

For categorical models, we observe that both Foldseek and the Categorical SoftAlign (20 characters) are based on categorical representations. However, the Categorical SoftAlign (20 characters) achieves an LDDT score of 0.53, outperforming Foldseek’s best configuration, which achieves an LDDT score of 0.47.

Compared to PLMAlign [18], a language-based method that uses sequence information to predict alignments, the SoftAlign model demonstrates substantial improvements in alignment quality. PLMA-lign achieves an LDDT score of 0.47, which is significantly lower than the SoftAlign score, with a gain of 0.09 in LDDT. This performance gap highlights the advantage of SoftAlign’s structural modeling, which directly incorporates spatial and structural features into its alignment process, enabling it to capture the nuances of protein geometry more effectively than PLMAlign’s sequence-focused approach. These results underline the importance of utilizing structural information for achieving higher-quality alignments in challenging test scenarios.

The softmax-based variants of SoftAlign exhibit comparatively lower performance. This outcome is expected, as these variants rely on a pseudo-alignment approach making them inherently less robust compared to real alignment methods. However, these models can be enhanced at inference time by applying the Smith-Waterman algorithm to enforce sequential alignments, effectively rescuing their performance.

In addition, the scatter-plots in Figure 3 illustrate the alignment scores between our Smith-Waterman SoftAlign model and TM-align/Foldseek. Figure 1A depicts the scatter plot of LDDT scores between our SoftAlign model and TM-align, while Figure 1B illustrates the corresponding scores between SoftAlign and Foldseek. In conclusion, our SoftAlign model demonstrates competitive performance compared to the state-of-the-art aligner TM-align, and outperforms other aligners, as evidenced by both quantitative metrics in Table 1 and visual representations in Figure 3.

**Figure 3.**
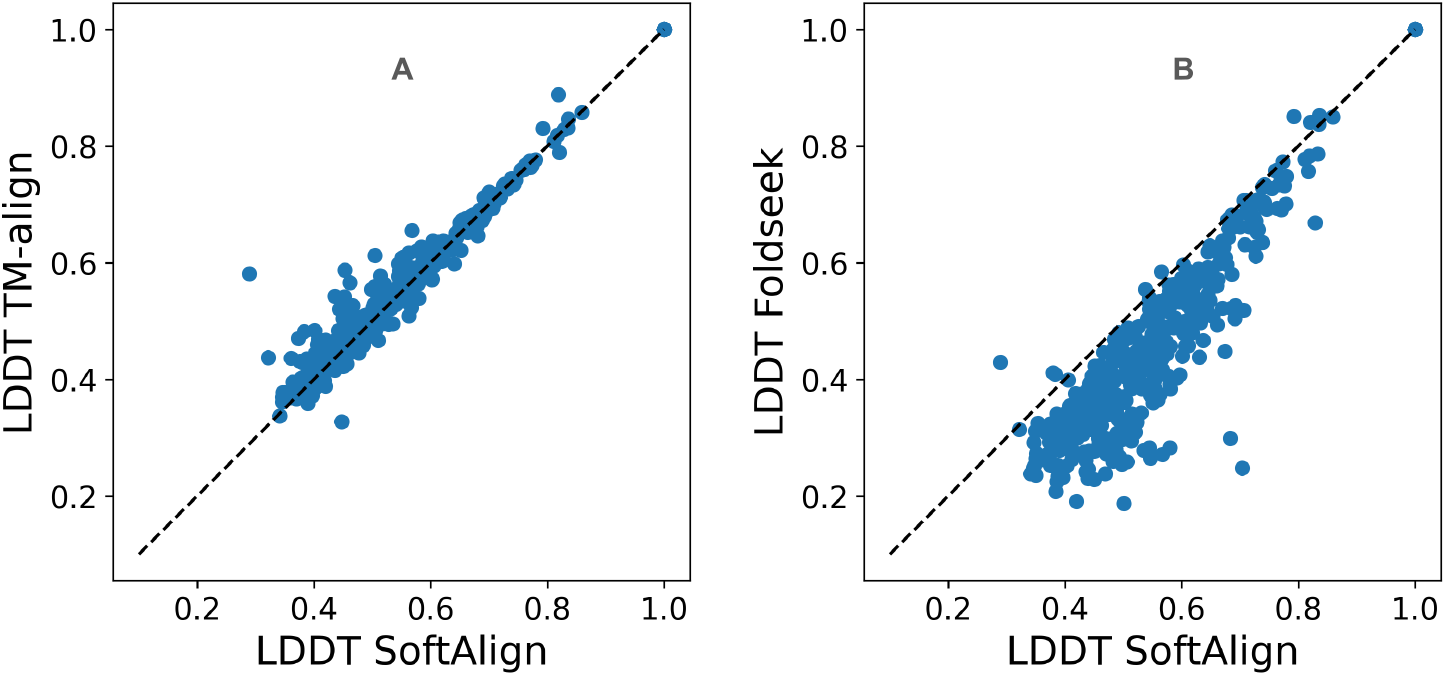
(a): Scatter plot comparing LDDT scores obtained from the SoftAlign model (x-axis) and TM-align (y-axis). (b): Same comparison using Foldseek instead of SoftAlign. The evaluation is performed on a test set of 1000 protein pairs from unseen folds, with TM-score *>* 0.6.

## Structure search

We measured the sensitivity of our model on the SCOPe40 dataset [13], where the ranking of alignments is determined by LDDT values. In contrast, Foldseek [14] uses a combination of TM-score and LDDT for ranking alignments. It is worth noting that we did not optimize for this ranking metric, as our primary focus was on alignment quality itself. However, integrating additional ranking metrics, such as the combination used by Foldseek, could potentially improve the performance of our model in future work. We used the same procedure as [14], performing an all-versus-all search and computing the performance in identifying members of the same family, super-family, and fold. The performance is measured by the fraction of true positives out of all possible correct matches until the first false positive, where a false positive is defined as a match to a different fold. The average is computed over all test proteins in the dataset. We compare the performance of our non-categorical and categorical SoftAlign models to the sensitivity of Foldseek, TM-align, and DALI.

Based on the results in Table 2, our non-categorical SoftAlign softmax model outperforms all structural aligners, including DALI. Additionally, the non-categorical and categorical SoftAlign models using the Smith-Waterman algorithm (SW) achieve competitive results compared to other methods.

**Table 2:** Sensitivity on the SCOPe40 dataset, comparison between Foldseek, TM-align, DALI and our SoftAlign (Sfa) models

| Model | Family | Super-family | Fold |
| --- | --- | --- | --- |
| Foldseek 3Di | 0.84 | 0.47 | 0.11 |
| Foldseek 3Di+sequence | 0.86 | 0.49 | 0.11 |
| TM-Align | 0.89 | 0.56 | 0.17 |
| TM-Align-fast | 0.89 | 0.56 | 0.16 |
| DALI | 0.89 | 0.58 | 0.17 |
| **Sfa Softmax** | **0.90** | **0.58** | **0.17** |
| Sfa SW | 0.86 | 0.53 | 0.15 |
| Sfa Categorical | 0.82 | 0.47 | 0.12 |

When comparing our categorical SoftAlign model with Foldseek, we observe no improvement. In fact, the performance of our categorical model is comparable to that of Foldseek, and there is no clear advantage to using our structural alphabet over Foldseek’s existing alignment methods. Rather than offering substantial benefits in this case, our findings highlight the need to bridge the gap between categorical and continuous methods, exploring ways to optimize categorical approaches for improved performance.

In the Discussion Section, we provide potential explanations for the effectiveness of the softmax approach. We also want to emphasize that the softmax method is not specific to our architecture and can be applied with other embedding methods, showcasing its flexibility and broad applicability across different alignment frameworks.

## Discussion

### End-to-end alignment

We introduce SoftAlign, a novel method designed to perform end-to-end alignment by directly mapping a pair of protein structures to an alignment. The combination of structural information as input and fully end-to-end training is essential for achieving near state-of-the-art alignment quality. SoftAlign is trained using two key loss functions:

LDDT Loss: We developed a differentiable version of the LDDT (Local Distance Difference Test) loss (see Methods section), making it suitable for gradient-based optimization. Unlike standard LDDT, which is non-differentiable, our smoothed formulation allows SoftAlign to directly optimize structural similarity during training. While this approach provides a self-contained alignment objective, further refinements are needed to improve local alignment accuracy.

Cross-Entropy Loss: The current implementation of SoftAlign, trained with cross-entropy loss, demonstrates competitive performance as an aligner. It surpasses other deep learning-based alignment methods in terms of alignment quality, showcasing the necessity of end-to-end training.

Although SoftAlign was specifically optimized for alignment tasks rather than structure search, it achieves the best performance for structure search as well. This suggests that the method could be further enhanced by explicitly incorporating structure search objectives into the training process. For instance, integrating a contrastive learning framework into the loss function could boost its effectiveness for structure search applications.

### Softmax: a good discriminator but a bad aligner

#### Lower Global LDDT and TM Scores

In our evaluations, as seen in Table 1, we observed that the softmax-based alignment method consistently produced lower global LDDT scores compared to traditional alignment methods like Smith-Waterman. The lack of sequential constraints in the softmax approach leads to an alignment that does not accurately capture the overall structural correspondence between proteins. However, as demonstrated in Table 1, this limitation can be rescued by applying the Smith-Waterman algorithm at inference time, which improves overall performance.

#### Effective Discrimination of False Positives

Despite its shortcomings as an aligner, the softmax method performs well in discriminating true positives from false positives in the structure search. False positives generated by the softmax method tend to cluster around LDDT scores close to zero, indicating that the method can effectively identify mismatches with low alignment quality. This behavior contrasts with the Smith-Waterman method, where false positives tend to have higher and more variable LDDT scores. This suggests that the Smith-Waterman algorithm, may sometimes generate alignments that appear superficially accurate, even when they do not reflect the true structural correspondence. In contrast, the softmax method, which does not rely on forcing an alignment path, can essentially align nothing if no meaningful structural correspondence exists, making it more conservative in its predictions and potentially better at avoiding false positives in cases of poor alignment.

Figure 4 provides a comparative analysis of LDDT values across the two different alignment methods. Panel (a) presents a scatter-plot for the query “d1kf6b2,” comparing LDDT scores from the softmax-based alignment (x-axis) and the Smith-Waterman method (y-axis). False positives (proteins from different folds) are shown in black, illustrating how the softmax method tightly clusters them around zero, whereas the Smith-Waterman method assigns some false positives relatively high scores. Panel (d) shows histograms of Z-scores of LDDT scores for all protein pairs in the SCOPe40 dataset. Each panel contains histograms for both the Smith-Waterman (SW) method and the softmax method. For the softmax method, Z-scores are higher when structural similarity is present (family, superfamily, and fold). In contrast, Z-scores for different folds are more concentrated around zero. This demonstrates that the softmax method is more effective at differentiating false positives from true structural similarities compared to the Smith-Waterman method.

**Figure 4.**
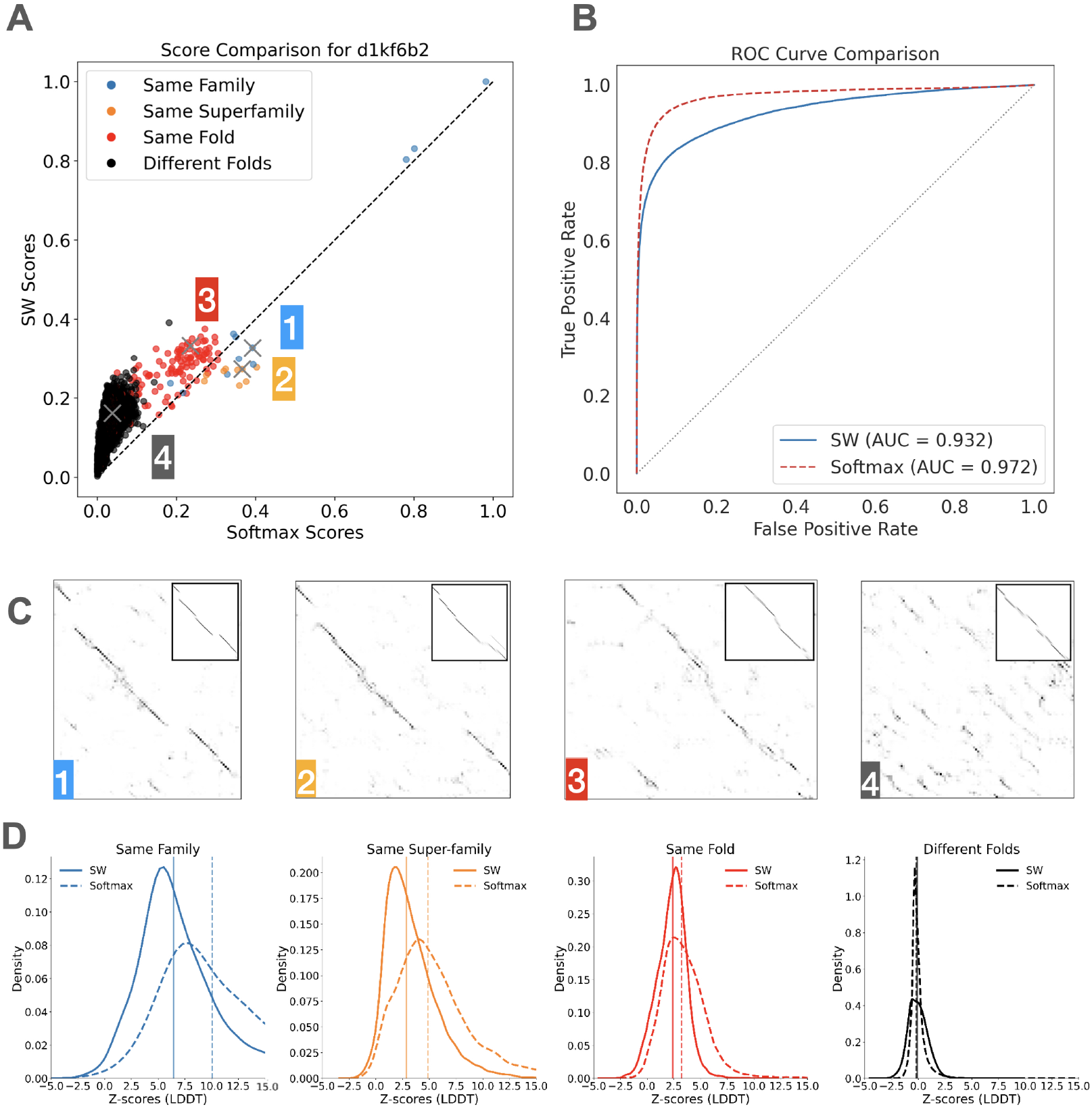
(a): Scatter plot of LDDT values comparing the softmax-based alignment method (x-axis) and the Smith-Waterman method (y-axis) for the query “d1kf6b2.” Black dots represent proteins belonging to a different fold (false positives).(b) ROC curve comparing the softmax-based alignment method and the Smith-Waterman method in distinguishing structurally related protein pairs. True positives include pairs from the same family, superfamily, or fold, while false positives are pairs from different folds. (c) Comparison of alignments produced by the softmax method and Smith-Waterman for the same query d1kf6b2 across different levels of structural similarity: family (1), superfamily (2), fold (3), and non-fold matches (4). The main plots correspond to the softmax-based alignments, while the Smith-Waterman alignments are insets within the plots. The softmax method produces alignment-like maps for family, superfamily, and fold levels but fails to generate meaningful alignments at the non-fold level, resulting in sparse and scattered points. In contrast, the Smith-Waterman method creates alignments at all levels, even for non-fold matches, due to its sequential constraints, resulting in forced alignments that may misrepresent the lack of true structural similarity.(d): Histogram of Z-scores of LDDT scores for all protein pairs in the all-vs-all search on SCOPe40 for both the softmax-based and Smith-Waterman alignment methods at the family, super-family, fold, and different folds level.

#### Impact of Non-sequential Property

The superior discrimination capability of the softmax method can be attributed to its non-sequential nature. By not enforcing sequential alignment, the softmax method aligns only those positions that exhibit strong mutual agreement in similarity. This is evident when computing the LDDT/TM-scores per aligned position instead of normalizing it by the number of positions in the query; only positions with high similarity, as indicated by both row-wise and column-wise softmax, are aligned. Consequently, fewer positions are aligned overall, but these positions are more reliably similar. Table 3 shows that while the softmax peudo-alignments are significantly worse than TM-align and Smooth Smith-Waterman alignments when the scores are normalized by the query length, they are of the same quality when the scores are normalized by the number of aligned positions.

**Table 3:**
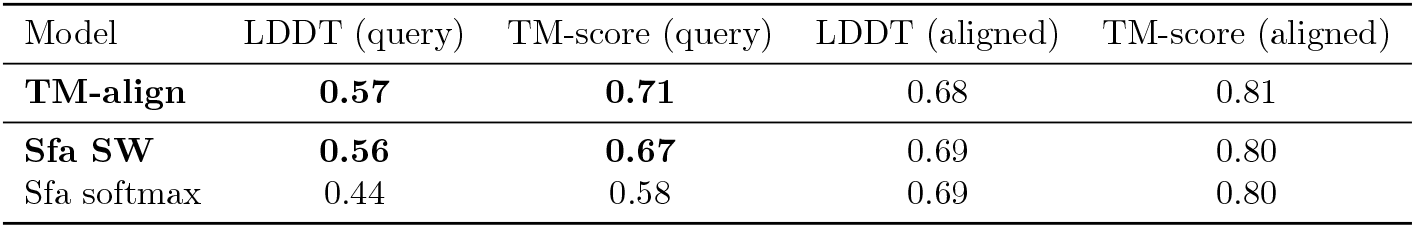
Comparison of LDDT and TM-scores for different alignment methods, normalized either by the query length (LDDT/TM-score query) or by the number of aligned positions (LDDT/TM-score aligned). While the softmax-based pseudo-alignments yield lower scores when normalized by query length, their scores are comparable to TM-align and Smooth Smith-Waterman when normalized by the number of aligned positions, indicating high reliability in the aligned regions.

#### Characteristics of False Positive Alignments

As shown in panel (c) of Figure 4, the alignment maps illustrate four representative cases across increasing structural divergence: same family, same superfamily, same fold, and different folds (false positives). Each case displays both the softmax alignment and, in the inset, the corresponding Smith-Waterman alignment. In the false positive scenario (panel 4), the softmax alignment fails to form a coherent alignment path—instead, only a few scattered high-similarity points appear in the map, reflecting low structural similarity and resulting in a very low global LDDT score. This sparsity highlights the softmax model’s ability to effectively reject structurally mismatched inputs. In contrast, the Smith-Waterman method, due to its sequential constraints, produces an alignment even in this mismatched case, leading to a more continuous but misleadingly structured path that does not reflect true correspondence. While Smith-Waterman is theoretically capable of skipping alignment altogether, in practice, it tends to force alignments,resulting in higher LDDT variability for false positives.

In summary, while the softmax method is not suitable as a primary alignment tool due to its lower global LDDT and TM scores, it demonstrates a very good capability as a discriminator of true and false positives. Its non-sequential nature allows it to focus on highly similar positions, resulting in effective filtering of false positives. This makes the softmax method a valuable complementary tool in protein structure search, particularly in scenarios where distinguishing low similarity from noise is critical.

### Categorical embeddings

#### Proposed categorical version of SoftAlign

To explore an alternative to Foldseek’s predefined alphabet, we proposed a categorical version of our SoftAlign framework. This model produces embeddings represented as discrete cluster identifiers, forming an alternative alphabet tailored to the data. While the categorical SoftAlign shows notable improvements in alignment quality compared to Fold-seek, particularly in terms of global LDDT and TM-scores, it does not provide similar enhancements for structure search tasks.

### The softmax method is not compatible with categorical values

One of the limitations of the softmax alignment method comes from its inability to effectively handle the discrete nature of embeddings in the categorical SoftAlign. The softmax alignment strategy aims to assign probabilities to each pair of positions along the rows and columns of the similarity matrix. However, a critical challenge happens when identical values are present in the similarity matrix due to the discrete nature of embeddings. In the categorical SoftAlign framework, positions are associated with discrete identifiers representing cluster centers. As a result, the similarity matrix contains a finite set of possible values determined by the number of clusters. More precisely, there are exactly 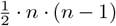 possible values, where *n* is the number of clusters. When softmax is applied to such similarity matrices, it struggles to differentiate between identical values. Since softmax assigns probabilities based on the relative magnitudes of values within each row or column, identical values yield identical probabilities after softmax transformation. Consequently, softmax alignment fails to offer meaningful distinctions among position pairs with identical similarity scores.

### Runtime

We evaluated the runtime performance of SoftAlign (Softmax version) in an all-vs-all structural search on the SCOPe40 dataset, which consists of 11211 proteins. The total runtime for SoftAlign was approximately 6425 seconds, including both the alignment computations and embedding generation. For the embedding generation step, SoftAlign required around 3 minutes (∼ 180 seconds) to process the 11211 proteins.

For comparison, the runtimes of other methods are summarized in Table 4, measured in seconds for the same task.

**Table 4:** Comparison of runtime performance across different structural alignment methods.

| Method | Runtime (seconds) |
| --- | --- |
| TM-align (64-core CPU) | 162240 |
| TM-align-fast (64-core CPU) | 62240 |
| DALI (64-core CPU) | 123010 |
| Foldseek-TM (64-core CPU) | 1889 |
| Foldseek (64-core CPU) | 42 |
| SoftAlign (Softmax) (single GPU) | 6425 |

These results demonstrate that SoftAlign achieves competitive runtime performance compared to traditional methods like TM-align and DALI. Although Foldseek and its variants are faster, they show lower alignment quality compared to SoftAlign, as described in the previous sections. This highlights the balanced trade-off between runtime and alignment accuracy achieved by SoftAlign.

## Conclusion

The SoftAlign framework presents a powerful end-to-end approach to protein structure alignment.The model achieves competitive alignment quality and demonstrates unique capabilities in structure search tasks.

The softmax alignment method, despite its limitations as a traditional aligner, excels in discriminating true positives from false positives in structure search. Its non-sequential nature allows it to selectively align only the most similar positions, resulting in robust performance in filtering tasks. However, its pseudo-alignment approach results in lower global LDDT and TM-scores, a limitation that can be mitigated by applying Smith-Waterman refinement at inference time. This demonstrates the softmax method’s utility as a complementary tool for precise pre-filtering in large-scale protein databases.

The categorical SoftAlign provides an alternative structural alphabet, similar to Foldseek’s approach, with promising alignment improvements. While its current performance in structure search remains comparable to Foldseek, making it less competitive as a drop-in replacement, it may be capturing different aspects of structural information. This suggests that it could outperform existing methods in specific contexts, warranting further investigation into its strengths and use cases. Further work is required to optimize the categorical approach for broader applications.

In summary, SoftAlign is a precise alignment framework that bridges the gap between efficient structure search and accurate alignment. Future work will focus on enhancing the categorical method, improving runtime performance, and exploring the softmax-based alignment method integration with protein language model embeddings.

## Methods

### ProteinMPNN

ProteinMPNN is a message passing neural network that aims to find an amino acid sequence that will fold into a given structure [10]. The network takes as inputs the 3D coordinates and computes the following information for each residue: (i) the distance between the *N, C*_*α*_, *C, O* and a virtual *C*_*β*_ atom, (ii) the *C*_*α*_ − *C*_*α*_ − *C*_*α*_ frame orientation and rotation, (iii) the backbone dihedral angles, and (iv) the distances to the 64 closest residues. The full network is composed of an encoder and a decoder with 3 layers each. For our task, we retrained the encoder to obtain an embedding of each position that contains relevant information about the structure, and we do not use the decoder. The downstream task in our study is to create embeddings that can be used for alignment and similarity assessments, which is different from the original ProteinMPNN task of predicting amino acid sequences from structural data.

### Alignment strategies

#### Smooth and Differentiable Smith-Waterman

We apply the smooth and differentiable Smith-Waterman algorithm as implemented in [12]. The Smith-Waterman algorithm is a dynamic programming algorithm that inputs two sequences and a matrix of alignment scores for all pairs of positions and returns the optimal (highest scoring) alignment between the pair of sequences. The smooth version instead returns a distribution over alignments where higher scoring alignments are more likely. (This is achieved by replacing the *max* function with *logsumexp*. See [12] for details.) The extent to which this distribution is concentrated on the optimal alignment is controlled by the temperature parameter. The smoothing operation makes the algorithm differentiable, and therefore applicable in neural network pipelines.

#### Softmax alignment

We tested another strategy to create a pseudo-alignment by applying the softmax function on the rows and columns of the similarity matrix. Let **A** be the similarity matrix and *T* a temperature parameter. The softmax function transforms a vector of real numbers into a probability distribution. In this context, it is applied to each row and column of **A**.

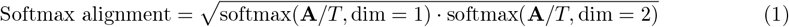

Focusing on the row-wise softmax application, for a specific position i in the first protein, the softmax values indicate the probability of i being structurally similar to each of the positions of the second protein. Positions with higher softmax scores are considered more likely to be structurally equivalent. Similarly, the column-wise softmax applied to a specific position j in the second protein creates a probability distribution. Here, each element represents the probability that position j aligns well with the corresponding position i in the first protein.

The product of the row-wise and column-wise softmax outputs is taken because we want to capture agreement between the two perspectives. Elements with high probability in both their corresponding row and column will contribute more significantly to the final result.

Note that this method, unlike the Smith-Waterman aligner, does not impose a sequential alignment. If the position *i*_1_ in the first protein is aligned with *j*_1_ in the second, and *i*_2_ with *j*_2_, then if *i*_1_ *< i*_2_, the softmax method does not guarantee *j*_1_ *< j*_2_. In the discussion section, we will address the drawbacks and advantages of this method. Figure. 2 provides an example of the two alignment methods.

### Architecture of the SoftAlign alignment network

The network, called SoftAlign, takes as input the 3D coordinates of the two proteins. These coordinates are transformed into a vector of features for each position using the ProteinMPNN architecture. Note that we do not use the weights of ProteinMPNN; we instead trained the ProteinMPNN encoder using a structural alignment task. The scalar products are taken between the feature vectors of each the two proteins to obtain a similarity matrix *A* with matrix elements *a*_*i,j*_ corresponding to the score of aligning *i* with *j*. Lastly, the Smooth-Smith-Waterman algorithm, or the softmax strategy is applied to the similarity matrix to obtain an alignment. The Smooth-Smith-Waterman and the softmax method algorithms outputs the probability that a position pair is aligned, therefore we obtain a soft alignment. In order to get an hard alignment, in other words to have **P**(i,j aligned) = 0 or **P**(i,j aligned) = 1, we take a very low temperature *T* in the scoring function. The temperature is therefore decreased during training from *T* = 5 to *T* = 1. At inference time, we set *T* to 10^−4^. This workflow is summarized in Figure. 1.

### Categorical SoftAlign

To enable the use of sequence-based preprocessing tools for database filtering [16], we introduce a discretization step for both the query and target structures. To achieve this, we developed Categorical SoftAlign, a modified version of our SoftAlign network that incorporates discretization.

### Discretization via Clustering

Instead of representing each sequence position with a continuous vector (i.e., the output of the MPNN encoder), we assign each position a discrete identifier corresponding to a cluster center. The process unfolds as follows:

1. **Initial Training**: We first train the network using the standard SoftAlign approach.
2. **Cluster Initialization**: We embed all protein structures from the training set and apply *k-means clustering* to determine initial cluster centers.
3. **Continued Training with Cluster Updates**: After obtaining the initial centers, we resume training, allowing the cluster centers to be updated dynamically.

### Obtaining Discrete Identifiers

To assign a categorical variable to each position during training, a straightforward approach would be selecting the identifier of the nearest cluster center. However, this operation is not differentiable. Instead, we use the following differentiable approximation:

1. Compute the dot product between the initial embeddings and the cluster centers:

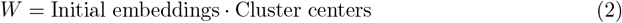
2. Construct a membership vector for each position using the *argmax operation* with the straight-through trick:

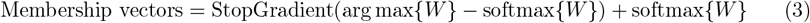

This ensures that during backpropagation, gradients flow through the softmax while maintaining a discrete selection.
3. Obtain the final discretized embeddings by multiplying the membership vectors with the transposed cluster center matrix. This step maps each position to a learned cluster center:

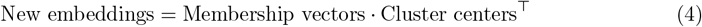

### Importance of k-Means Initialization

The initialization of cluster centers via k-means is crucial. We observed that without this step, the network tends to get stuck in local minima, leading to poor clustering performance. By contrast, initializing with k-means provides a strong starting point for meaningful cluster assignments. This entire discretization process is summarized in Figure. 5.

**Figure 5.**
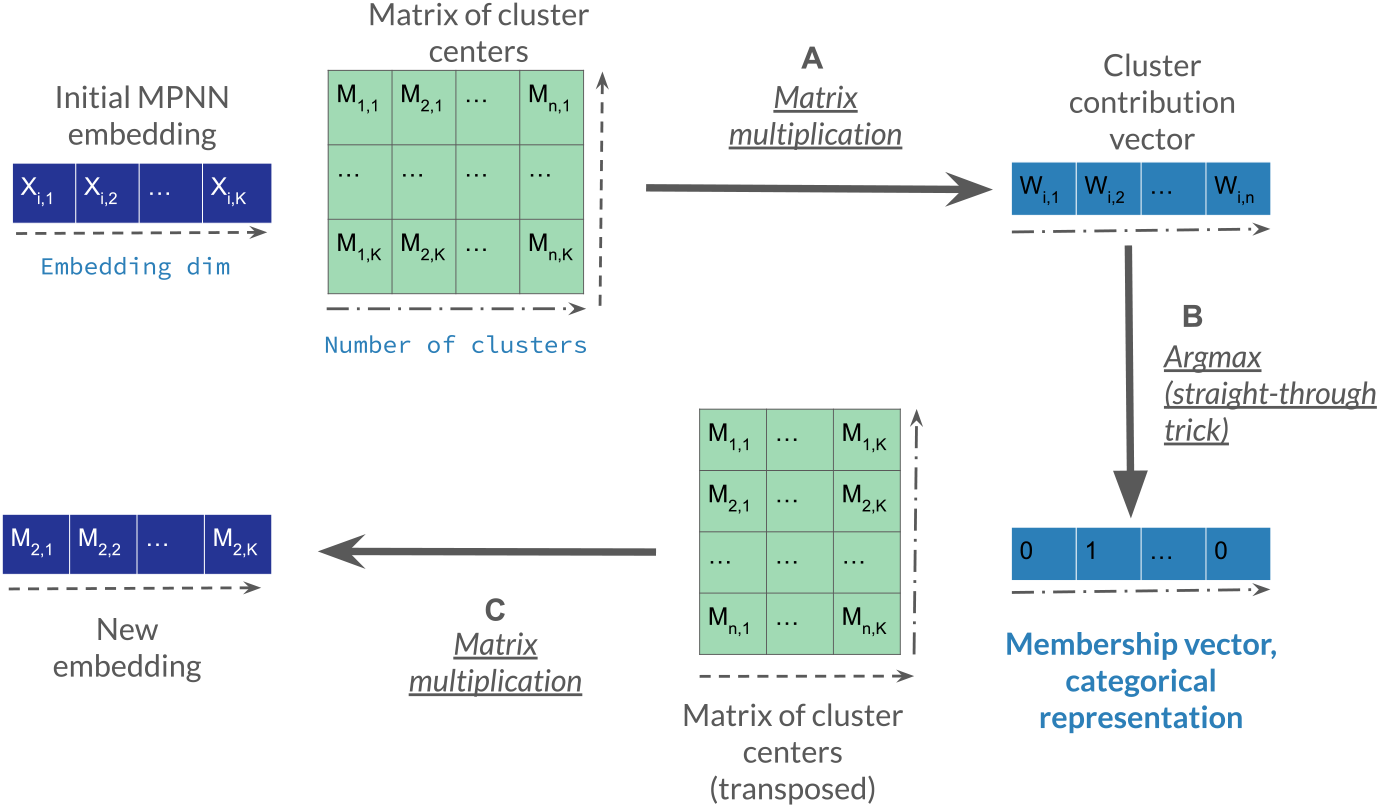
Discretization step of Categorical SoftAlign. (a) The MPNN embedding at a given sequence position is multiplied by the matrix of cluster centers, producing a vector where each element *i* represents the contribution of cluster *i*. (b) The argmax operation is applied to obtain the membership vector, a one-hot encoding of the most relevant cluster. (c) The membership vector is then multiplied by the transposed cluster center matrix to reconstruct an embedding corresponding to the assigned cluster center.

### Loss Functions

We experimented with two types of loss functions to train the network: cross-entropy loss and a smoothed LDDT loss.

### Cross-Entropy Loss

The first loss function we consider is the cross-entropy (CE) between the structural alignment predicted by TM-align and the alignment generated by our model:

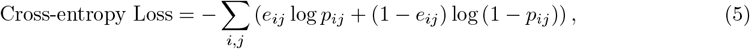

where - *e*_*ij*_ = 1 if TM-align aligns position *i* in the first protein with position *j* in the second protein, and 0 otherwise, - *p*_*ij*_ is the predicted probability that the two positions are aligned according to our method.

## Smoothed LDDT Loss

The second loss function is a smoothed version of the local distance difference test (LDDT) score, which evaluates how well the local atomic environment in a reference structure is preserved in the aligned structure.

A key advantage of this loss function is that it does not rely on external computations such as TM-align, making our model more end-to-end. Unlike TM-score or GDT-score, LDDT is also invariant to the relative orientation of domains, making it more robust for flexible proteins.

To compute the LDDT loss:

1. We first compute the pairwise distances between all *C*_*α*_ atoms in the reference structure that are within a 15 ^°^A cutoff. This results in a pairwise distance matrix *D*_1_ of size *L*_1_ × *L*_1_, where *L*_1_ is the length of the query protein.
2. Similarly, we compute a pairwise distance matrix *D*_2_ for the aligned target structure. *D*_2_ has dimensions *L*_2_ × *L*_2_.
3. To correctly compare these matrices, we use the alignment matrix *P* (size *L*_1_ × *L*_2_) to project *D*_2_ into the query space:

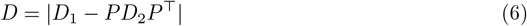
4. A pairwise distance in the aligned structure is considered conserved if its deviation is within a threshold *t*, i.e., if *D*_*i,j*_ < *t*. Instead of using a hard threshold, we smooth the decision using a sigmoid function to make it differentiable:

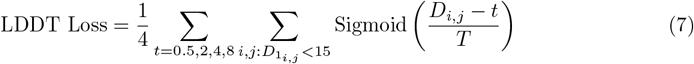

The temperature parameter *T* controls the sharpness of the sigmoid function. In the limit *T* → 0, we recover the traditional LDDT score. During training, *T* was progressively reduced from 5 to 0.1 to allow for a smoother optimization process.

## Effect on Alignment Strategy

Our LDDT-based loss is normalized by the query length rather than the number of aligned positions. This prevents the model from focusing on aligning only a small number of highly accurate positions and instead encourages global alignment.

The results of this approach are detailed in Section as a proof of concept. However, for practical applications, additional improvements are needed to extend the method to effectively support local alignments and use it for structure search.

## Training set

The SCOPe40 [13] dataset, containing 11,211 protein domains clustered at 40% identity, was chosen for its focus on single domains with known structures. However, a standard split for train and test based on sequence identity might not be ideal. Due to the nature of proteins, it is possible for some protein pairs to share a low degree of sequence similarity (below 40% identity in this case) yet exhibit a high degree of structural similarity. Therefore, to ensure a more robust evaluation of model generalization and performance, we employed a split based on protein folds rather than solely relying on sequence identity. Our training set includes 6506 proteins clustered in 681 different folds. Our test set includes 4507 proteins clustered in 513 different folds. Since we train using cross-entropy loss, we selected protein pairs with a TM-score greater than 0.6, ensuring that the chosen pairs exhibit significant structural similarity and produce meaningful TM-align alignments.

## Acknowledgements

We thank the Agence National de Recherche (ANR) IHU FOReSIGHT (ANR18-IAHU-01) and the I-Demo Regional (BPI, Région Île-de-France) grant GEAR (n° 407307) for funding support. Jeanne Trinquier was supported by a PhD Fellowship of the i-Bio Initiative from the Idex Sorbonne University Alliance. Samantha Petti was supported by a Burroughs-Wellcome Fund Careers at the Scientific Interface (CASI) award. Sergey Ovchinnikov acknowledges funding from NIH grant DP5OD026389, NSF MCB2032259, and Amgen.

We also thank Martin Weigt, Francesco Zamponi, and Ulisse Ferrari for insightful discussions and valuable feedback.

