## Supplementary for "SoftAlign: End-to-end protein structures alignment"


### Supplementary results

#### TM-scores

We evaluated TM-scores on the same SCOPe40 [1] test set, using TM-align [2] as the reference. Table 1 shows that our Smith-Waterman SoftAlign (Sfa SW) model achieves a TM-score of 0.67, closely matching DALI (0.68) [3] and outperforming categorical [4] and sequence-based [5] methods. The SoftAlign categorical variant (20 clusters) also surpasses Foldseek [4], further demonstrating the robustness of our approach.

While SoftAlign performs well, it is important to note that TM-align achieves the highest score (0.71) but has an unfair advantage in this comparison since it is specifically optimized for the TM-score.

Softmax-based SoftAlign variants initially underperform, but applying Smith-Waterman post-processing improves their TM-scores significantly (e.g., from 0.58 to 0.63). Figure 1 further illustrates the strong correlation between SoftAlign and TM-align scores, reinforcing the model's effectiveness for structural alignment.

#### Structure search

In Figures [2-8], we provide additional examples of scatter plots comparing LDDT scores between the Softmax and Smith-Waterman (SW) alignment methods. Specifically, we present 9 examples per SCOPe class, illustrating how the alignment scores vary across different structural categories. These visualizations further highlight the differences between the two methods in capturing structural similarities.

### Architecture and Training Details

Our model utilizes the encoder from ProteinMPNN [6], which consists of 3 layers with an encoding dimension of 64. For alignment, we employ the smooth and differentiable Smith-Waterman algorithm [7] with an affine gap penalty and learnable extension and open penalties. The model is trained using a learning rate of $10^{-3}$ for 50 epochs, with a decreasing temperature schedule from 5 to 1. Training is conducted on the SCOPe dataset [1], following the train-test split described in the main text. The full training code is available at `https://github.com/jtrinquier/SoftAlign`.

| Model | TM-score |
|---|---|
| Foldseek 3DI | 0.58 |
| Foldseek 3DI + sequence | 0.57 |
| PLMAlign | 0.59 |
| TM-align | 0.71 |
| DALI | 0.68 |
| **Sfa SW** | 0.67 |
| Sfa Softmax | 0.58 |
| Sfa Softmax rescued | 0.63 |
| Sfa SW LDDT Loss | 0.66 |
| Categorical Sfa 20 clusters | 0.64 |

Table 1: Comparison of mean TM-scores across different alignment methods. TM-align and DALI serve as traditional structural alignment baselines, while Foldseek and PLMAlign represent alternative approaches. Our SoftAlign (Sfa) models achieve strong performance, with the Smith-Waterman SoftAlign (Sfa SW) reaching a TM-score of 0.67, close to DALI. The "rescued" Sfa Softmax variant applies Smith-Waterman at inference time, improving its performance.

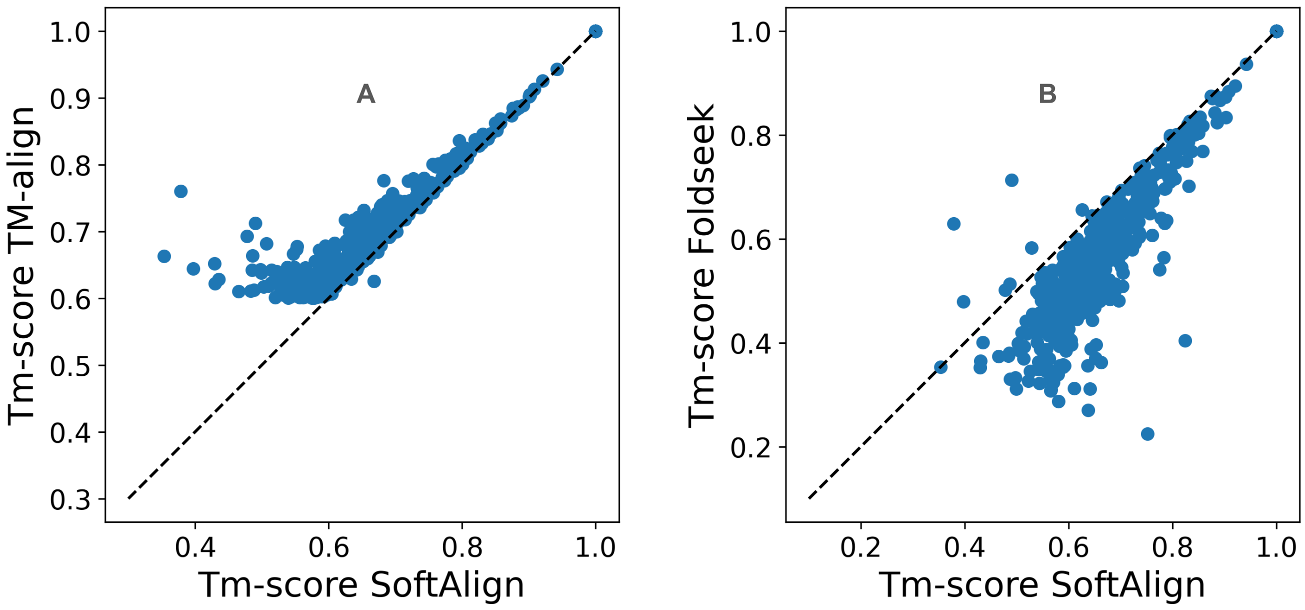


Figure 1: Figure A: Scatter plot of TM-scores between the SoftAlign-MPNN model (x-axis) and TM-align (y-axis). Figure B: Same comparison with Foldseek.

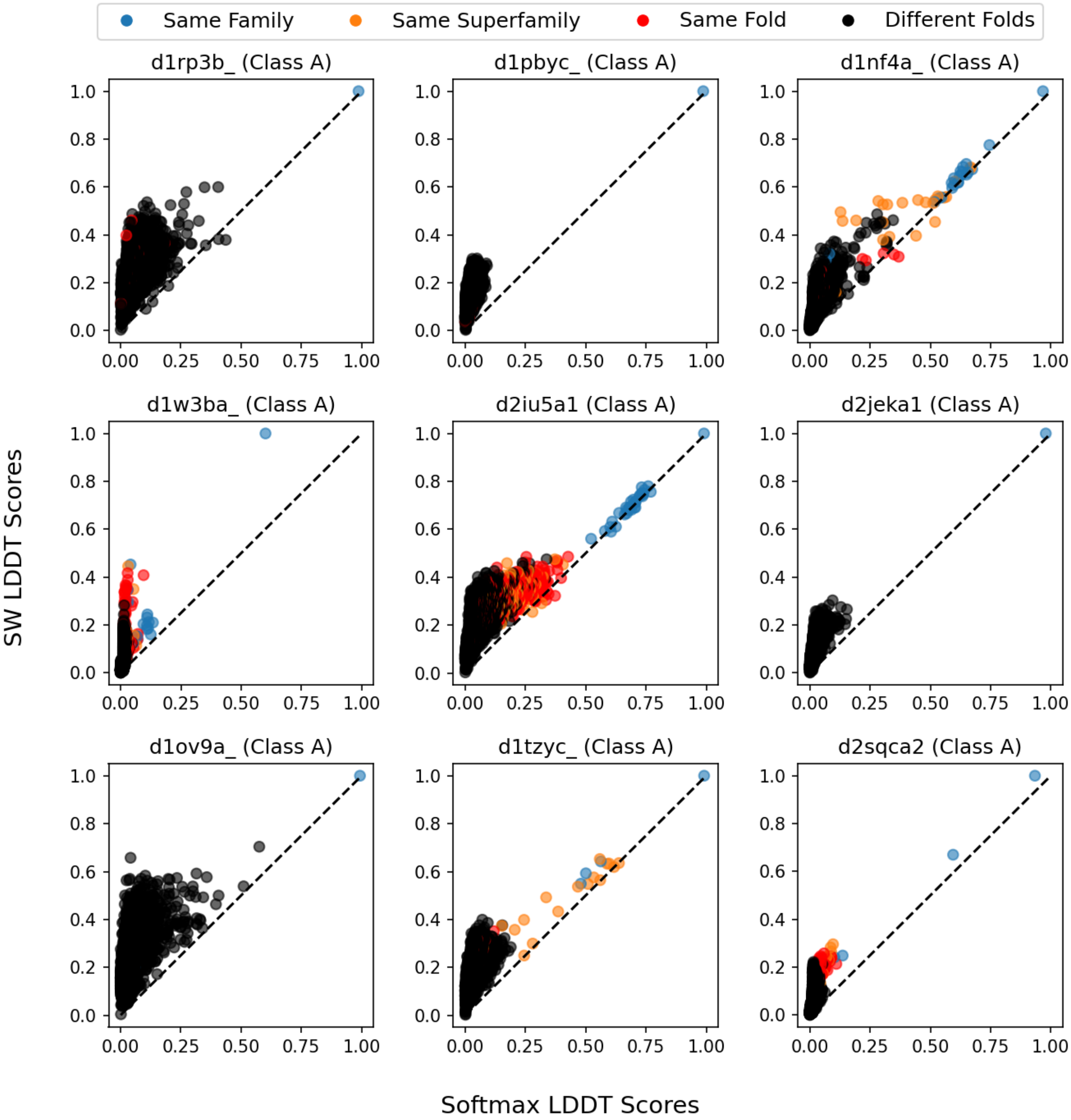


Figure 2: Scatter plot of LDDT values comparing the softmax-based alignment method (x-axis) and the Smith-Waterman method (y-axis) for eight queries in the Class A. Black dots represent proteins belonging to a different fold (false positives)

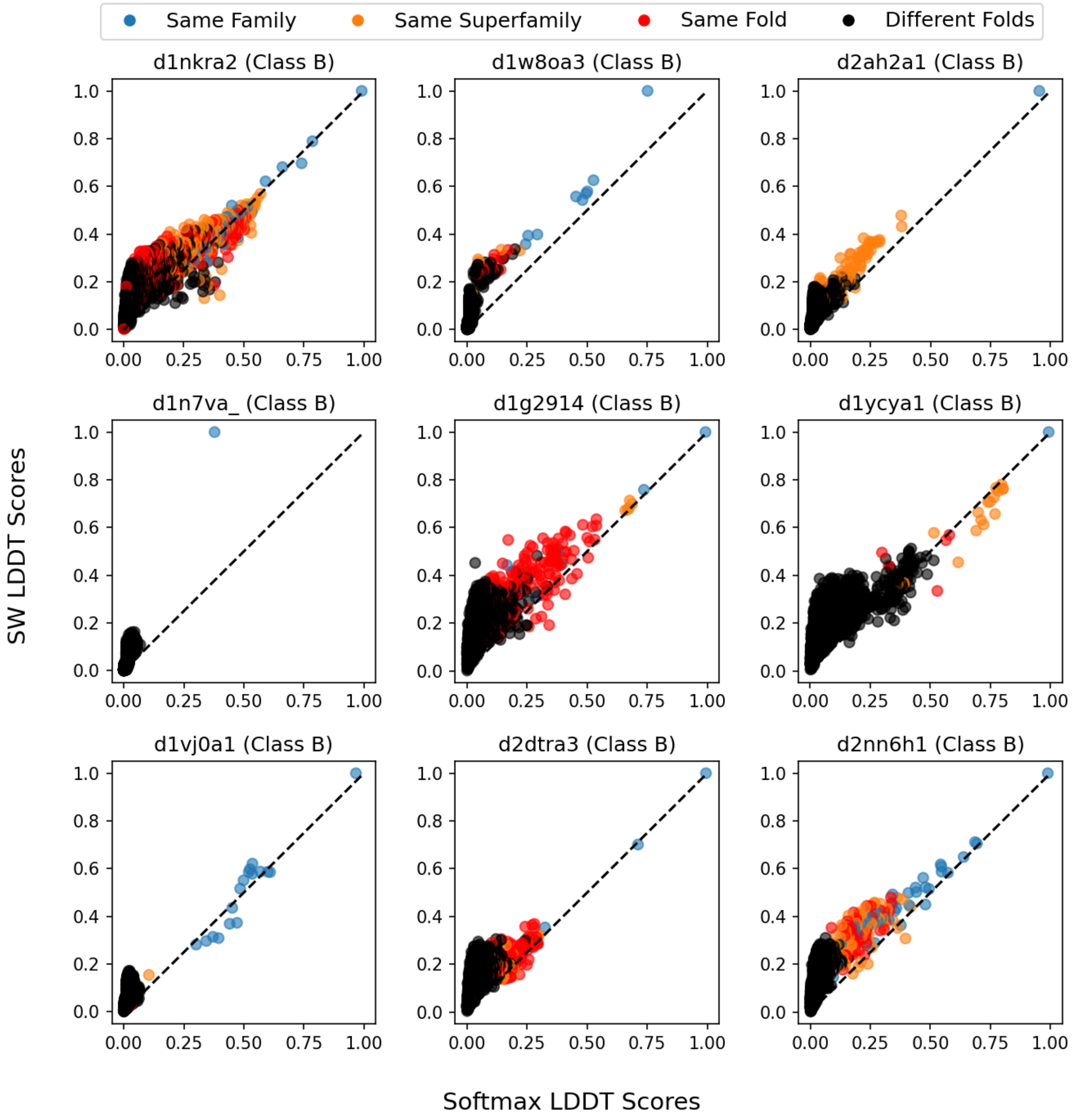


Figure 3: Scatter plot of LDDT values comparing the softmax-based alignment method (x-axis) and the Smith-Waterman method (y-axis) for eight queries in the Class B. Black dots represent proteins belonging to a different fold (false positives)

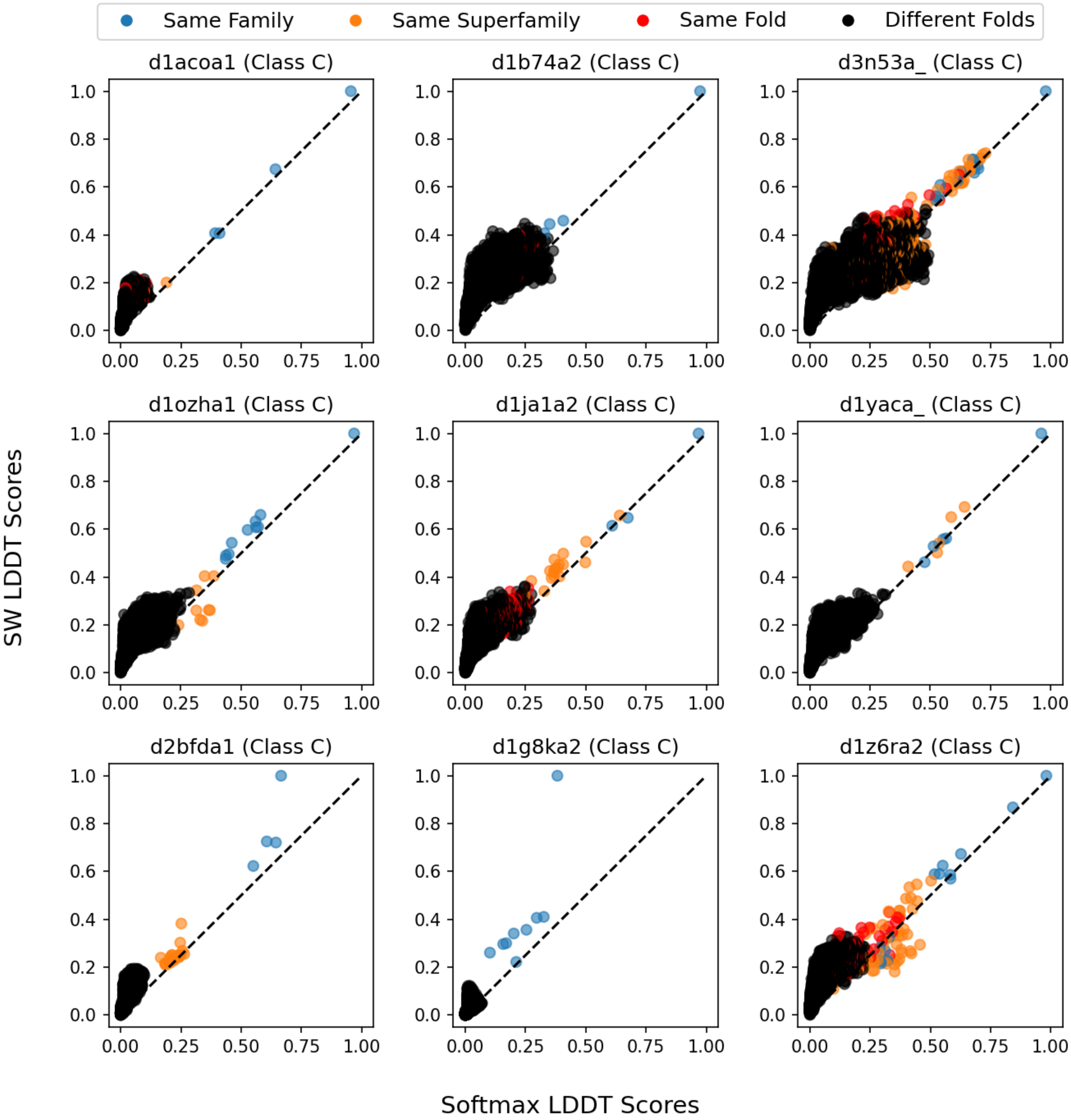


Figure 4: Scatter plot of LDDT values comparing the softmax-based alignment method (x-axis) and the Smith-Waterman method (y-axis) for eight queries in the Class C. Black dots represent proteins belonging to a different fold (false positives)

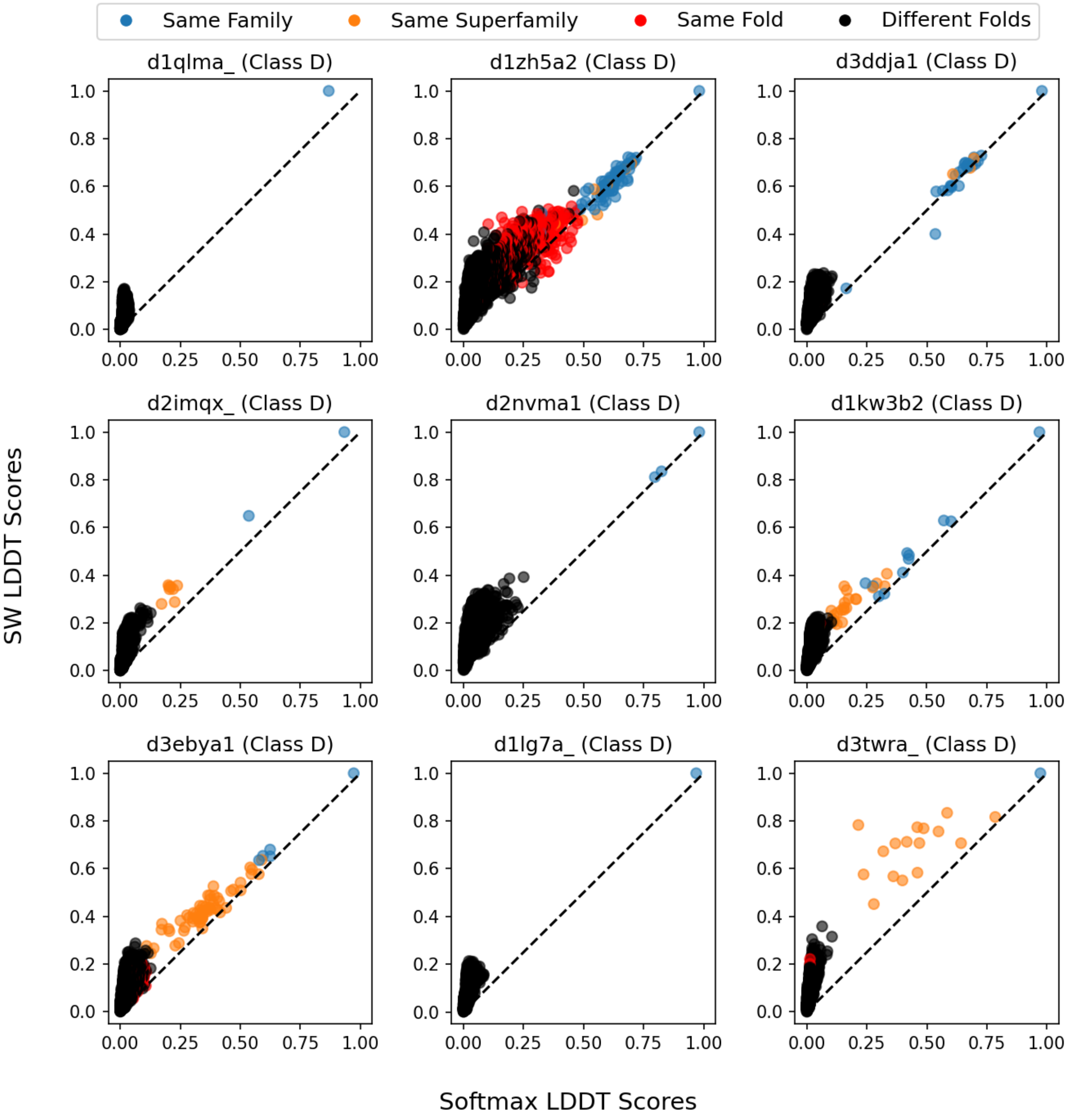


Figure 5: Scatter plot of LDDT values comparing the softmax-based alignment method (x-axis) and the Smith-Waterman method (y-axis) for eight queries in the Class D. Black dots represent proteins belonging to a different fold (false positives)

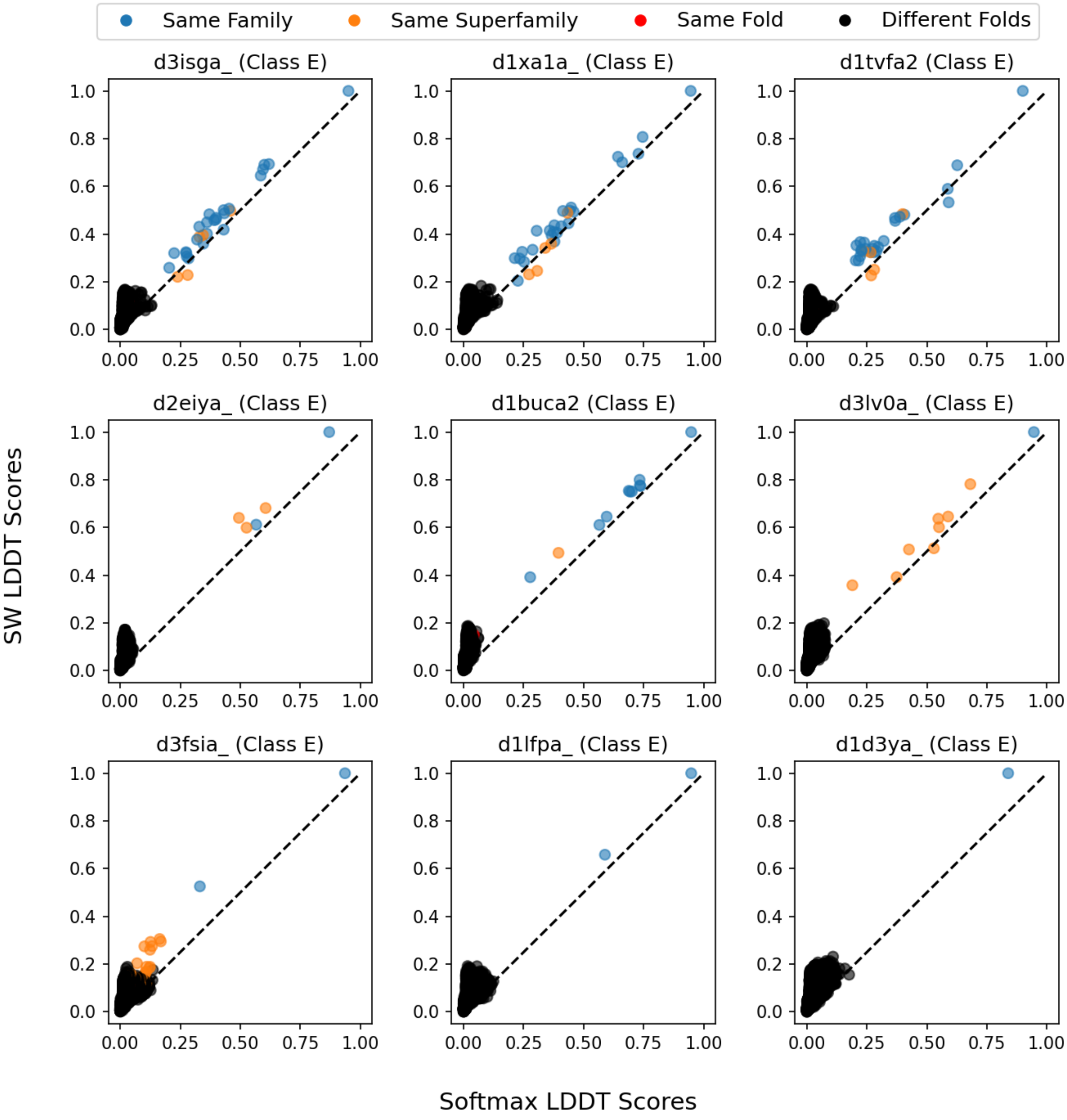


Figure 6: Scatter plot of LDDT values comparing the softmax-based alignment method (x-axis) and the Smith-Waterman method (y-axis) for eight queries in the Class E. Black dots represent proteins belonging to a different fold (false positives)

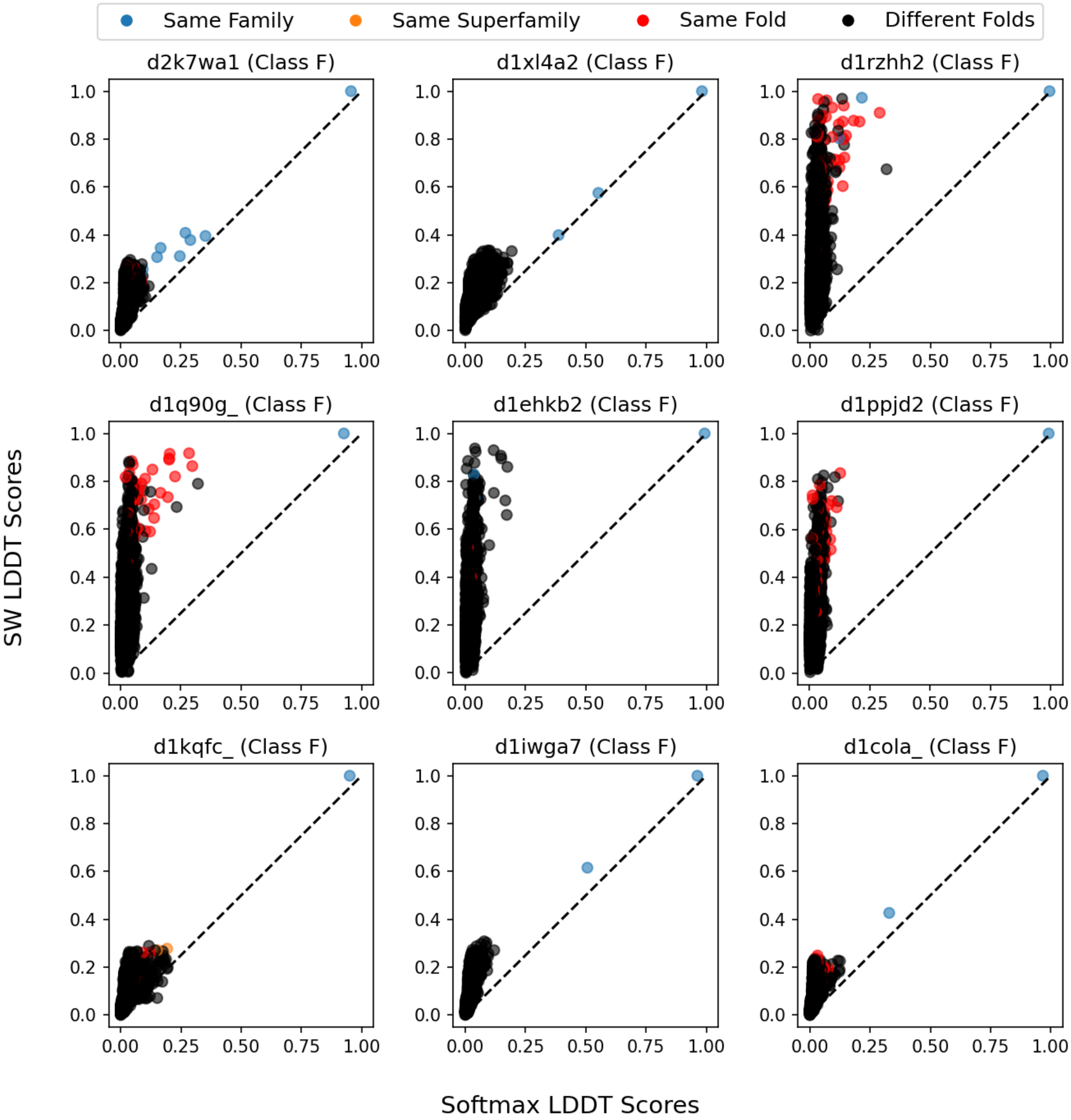


Figure 7: Scatter plot of LDDT values comparing the softmax-based alignment method (x-axis) and the Smith-Waterman method (y-axis) for eight queries in the Class F. Black dots represent proteins belonging to a different fold (false positives)

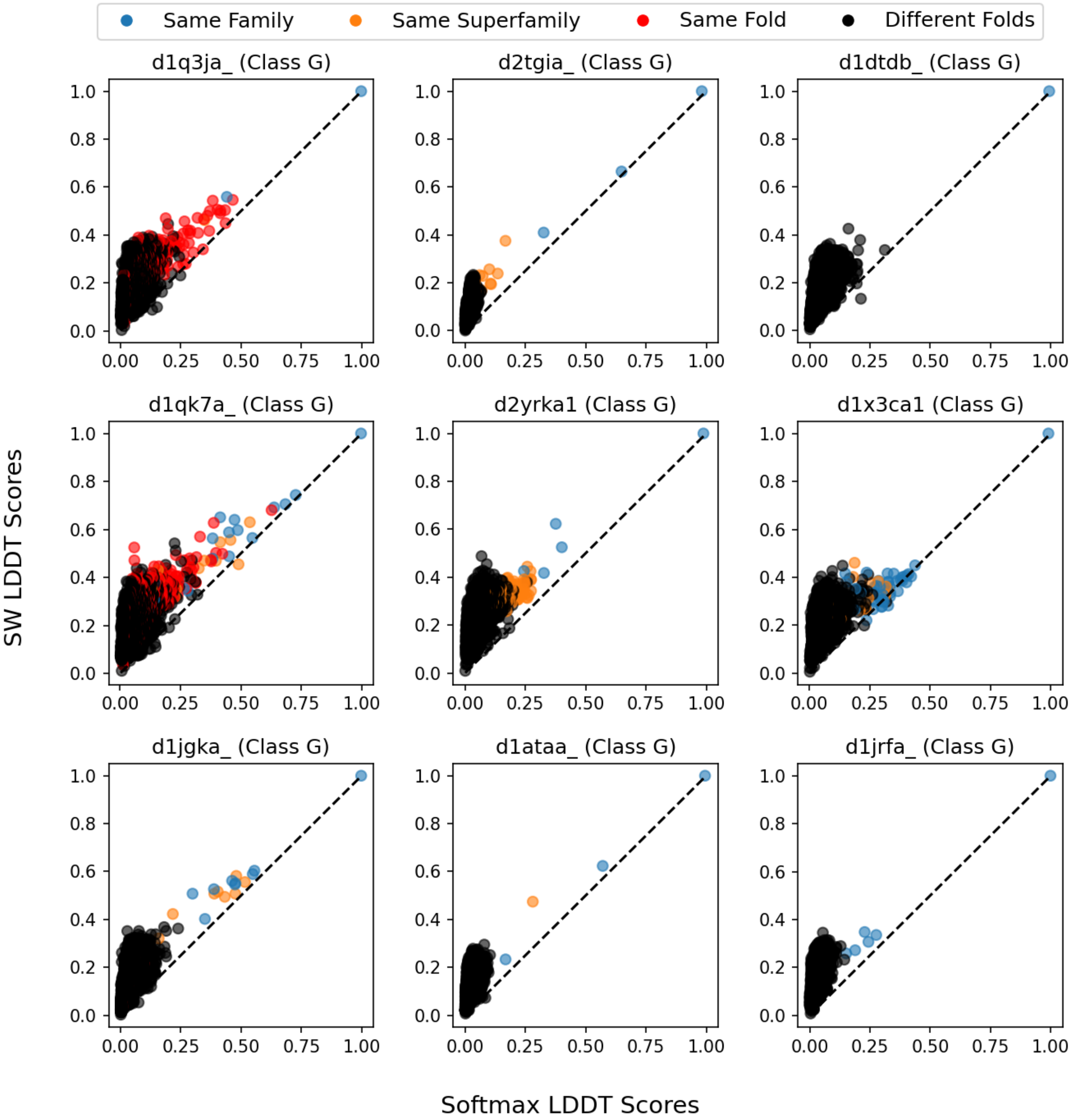


Figure 8: Scatter plot of LDDT values comparing the softmax-based alignment method (x-axis) and the Smith-Waterman method (y-axis) for eight queries in the Class G. Black dots represent proteins belonging to a different fold (false positives)
